# Trade-offs between cost of ingestion and rate of intake drive defensive toxin use

**DOI:** 10.1101/2021.07.23.453507

**Authors:** Tyler E. Douglas, Sofia G. Beskid, Callie E. Gernand, Brianna E. Nirtaut, Kristen E. Tamsil, Richard W. Fitch, Rebecca D. Tarvin

## Abstract

Animals that ingest toxins can become unpalatable and even toxic to predators and parasites through toxin sequestration. Because most animals rapidly eliminate toxins to survive their ingestion, it is unclear how populations transition from susceptibility and toxin elimination to tolerance and accumulation as chemical defense emerges. Studies of chemical defense have generally focused on species with active toxin sequestration and target-site insensitivity mutations or toxin-binding proteins that permit survival without necessitating toxin elimination. Here, we investigate whether animals that presumably rely on toxin elimination for survival can utilize ingested toxins for defense. We use the A4 and A3 *Drosophila melanogaster* fly strains from the Drosophila Synthetic Population Resource (DSPR), which respectively possess elevated and reduced metabolic nicotine resistance amongst DSPR fly lines. We find that ingesting nicotine increased A4 but not A3 fly survival against *Leptopilina heterotoma* wasp parasitism.Further, we find that despite possessing genetic variants that enhance toxin elimination, A4 flies accrued more nicotine than A3 individuals likely by consuming more media. Our results suggest that enhanced toxin metabolism can allow for greater toxin intake by offsetting the cost of toxin ingestion. Passive toxin accumulation that accompanies increased toxin intake may underlie the early origins of chemical defense.

## Introduction

Most animals survive toxin ingestion by eliminating toxins through metabolic detoxification (1–3). Some chemically defended animals subvert this paradigm by sequestering dietary toxins to deter predators or parasites (4). Because metabolic detoxification serves to prevent toxin accumulation, toxin-sequestering taxa often employ resistance mechanisms that do not degrade toxins (5). For example, target-site insensitivity (TSI), which results from mutations in a protein that prevent toxins from binding, is common in toxin-sequestering insects (6, 7). TSI sometimes co-occurs with toxin-binding proteins that scavenge toxins and prevent them from binding to targets (8–11). Such non-metabolic resistance mechanisms may facilitate the transition from toxin elimination to sequestration by decreasing reliance on toxin breakdown for survival (12).

Although metabolic detoxification degrades toxins, it is unclear whether reliance on this mechanism constrains chemical defense evolution. Metabolic detoxification permits toxin consumption and may ultimately lead to toxin sequestration so long as consumption outpaces degradation. To test this idea, we obtained two isofemale, homozygous strains of *Drosophila melanogaster* from the Drosophila Synthetic Population Resource (DSPR (13)) that possess high and low nicotine resistance (A3 and A4, Bloomington stocks 3852 and 3844), and exposed them to nicotine, a plant allelochemical that targets acetylcholine receptors (14). Although some drosophilids do feed on toxic food sources (15, 16) and the A4 fly strain may have experienced incidental nicotine exposure on tobacco farms that were prevalent at its collection site (17), drosophilids are not known to select nicotine-producing plants as hosts. Nevertheless, the genetic basis of nicotine resistance in *D. melanogaster* is extensively characterized, making this toxin well-suited to modelling the evolutionary origins of chemical defense (18). Compared to A3, A4 flies possess duplicate copies of cytochrome p450s *Cyp28d1* and *Cyp28d2* that are constitutively expressed at higher levels. A4 flies also overexpress the UDP-glucuronosyltransferase *Ugt86Dd*, while A3 harbors a mutation in this gene that significantly reduces nicotine resistance (19). *Ugt86Dd* is located in a Quantitative Trait Locus (QTL) that contributes 50.3% of the broad-sense heritability in nicotine resistance of DSPR lines, while a QTL containing *Cyp28d1* and *Cyp28d2* accounts for 5% (18, 20). The contributions of these three genes to nicotine resistance have been confirmed using gene knockout (21). Previous QTL and expression-QTL studies did not report evidence for TSI or toxin-binding proteins in A3 or A4 lines. While these mechanisms could exist, variation in metabolic enzymes appears to underlie the major difference between A3 and A4 nicotine resistance.

## Results and Discussion

We first quantified A3 and A4 nicotine resistance by estimating the median lethal concentration (LC50) of nicotine (Fig. 1). Because A4 flies had low viability in general, to compare LC50 between strains for this assay we normalized percent survival by the maximum survival of each line on control food (see supporting data for non-normalized values). The A4 LC50 was nearly twice that of A3 (LC50_A4_ = 1.9 ± 0.3 mM [mean ± SD], LC50_A3_ = 1.1 ± 0.2 mM; Fig. 1). While A3 survival decreased significantly at 0.5 mM nicotine, A4 survival was not significantly impacted until 1.75 mM. We proceeded to use an intermediate level of 1.25-mM nicotine for subsequent experiments.

**Figure 1.**
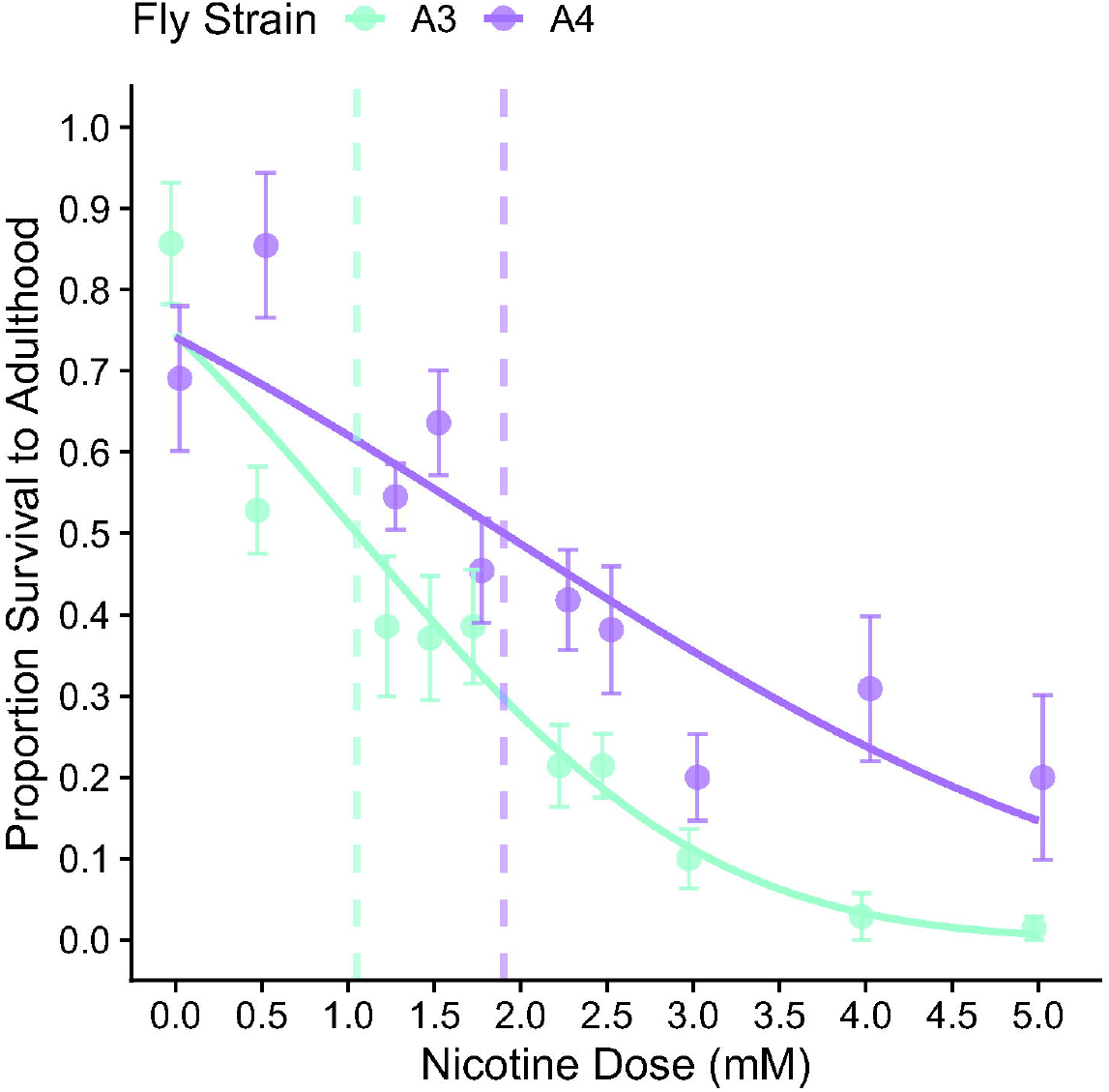
**A)** Nicotine concentration-survival curve for DSPR A3 and A4 *Drosophila melanogaster*. Data are normalized by maximum survival of each strain on control food. Vertical dashed lines represent LC50 of each strain.

We next assessed whether ingesting 1.25-mM nicotine after parasitism by the figitid wasp *Leptopilina heterotoma* increased *D. melanogaster* survival. *Leptopilina heterotoma* oviposits into the hemocoel of developing fly larvae, and actively suppresses the drosophilid defensive immune response against endoparasites (22). Thus, developing parasites are exposed to host hemolymph and, presumably, to circulating toxins consumed by fly larvae. In the control-fed, unparasitized treatment, 2.8% ± 2.7% of A4 larvae survived to adulthood, while in the nicotine-fed, parasitized treatment, A4 survival increased significantly to 6.8 ± 4.4% (p = 0.03, Z = -2.2; Fig. 2A). Correspondingly, *L. heterotoma* developmental success decreased five-fold from 37% ± 20% to 6.4% ± 6.8% (p < 0.0001, Z = 7.0; Fig. 2B). Thus, nicotine consumption increased A4 fly survival against parasitism.

**Figure 2.**
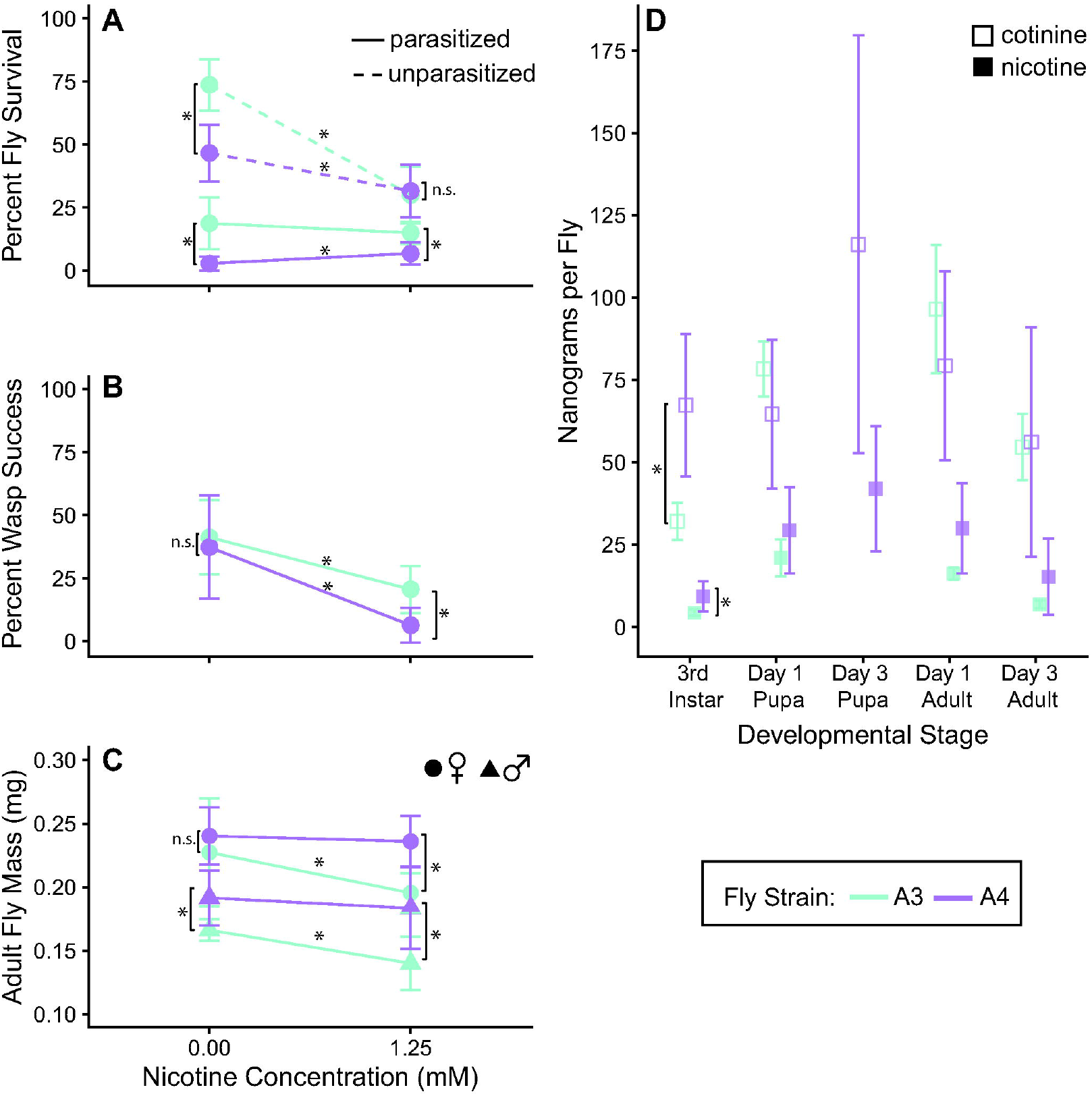
**A)** Nicotine consumption significantly decreases survival in unparasitized A3 and A4 *Drosophila melanogaster* flies. Nicotine consumption increases survival of parasitized A4 but not A3 flies. **B)** Nicotine consumption by A4 and A3 flies significantly decreases *Leptopilina heterotoma* developmental success. **C)** Nicotine consumption reduced A3 but not A4 adult body mass. **D)** Nicotine-fed A3 and A4 flies accumulate nicotine and its metabolic byproduct cotinine across developmental stages. Asterisks indicate significant differences.

In contrast, the survival of parasitized, nicotine-fed A3 larvae (15 ± 4.4%) was the same as parasitized, control-fed A3 larvae (19 ± 10%; p = 0.36, Z = 0.92; Fig. 2A). However, wasp developmental success on A3 flies halved from 41 ± 15% to 21 ± 9.3% when A3 flies consumed nicotine (p = 0.0001, Z = 4; Fig. 2B). This suggests nicotine consumption partially alleviated A3 parasitism-induced mortality. Nicotine consumption decreased unparasitized A3 fly survival by 44% (p < 0.0001, Z = 7.6), while nicotine consumption decreased parasitized A3 survival by only a tenth as much: 3.5%. The comparatively insignificant effect of nicotine consumption on parasitized A3 flies paired with a ∼50% decrease in wasp success suggests that nicotine may have offset parasitism-induced mortality for A3 flies, although to a lesser degree compared to A4 flies.

Next, we quantified nicotine accumulation in whole bodies of nicotine-fed larvae and adult flies. After 24hr ± 2.5hr on nicotine media, third-instar A4 larvae contained twice as much nicotine as A3 larvae (9.3 ± 4.6 vs. 4.3 ± 1.0 ng nicotine, p = 0.016, W = 1; Fig. 2D). Nicotine continued to accumulate until pupation and persisted through metamorphosis in both strains (Fig. 2D; also observed with ouabain [6]), suggesting that nicotine remained after the meconium was shed and may provide a defensive advantage into adulthood. The greater amount of nicotine in A4 could underlie the stronger effect of nicotine on parasite success in A4 versus A3 individuals (Fig. 2B). Although nicotine-fed A3 adults are ∼20% smaller than nicotine-fed A4 adults (Fig. 2C), this difference cannot explain the two-fold difference observed in nicotine accumulation between strains. The developmental rate of nicotine-fed A3 and A4 flies did not differ significantly at 1.25 mM nicotine and is also unlikely to underlie differences in nicotine accumulation (Fig. S1).

Our finding that A4 larvae accumulated more nicotine than A3 defies genotypic expectations, as A4 flies have genetic variants that are expected to increase nicotine breakdown (19, 21). To better understand this pattern, we compared relative amounts of cotinine, a metabolic by-product of nicotine (Fig 2D) between strains. A4 larvae contained significantly higher levels of cotinine compared to A3 individuals (Fig. 2D). Intriguingly, one-day-old and three-day-old A3 flies had significantly higher cotinine to nicotine ratios than A4, suggesting that A4 larvae have a distinct metabolic detoxification pathway compared to A3 (p_one-day-old_ = 0.031, W_one-day-old_ = 23, p_three-day-old_ = 0.008, W_three-day-old_= 25 (13)). This result matches expectations based on genotype, as the largest QTL underlying resistance in A4 contains several UGTs, which convert nicotine to glucuronides instead of cotinine (18).

The higher nicotine levels in A4 flies suggested that A3 flies are unable to survive high toxin loads, and thus might consume less to avoid nicotine accumulation. To quantify differences in feeding, we compared A3 and A4 adult body mass when reared on control versus nicotine food. While nicotine consumption significantly reduced A3 adult body mass, A4 mass remained unaffected (Fig. 2C), indicating that nicotine sensitivity constrained A3 food intake. The tobacco hornworm *Manduca sexta* employs a more extreme version of this pattern: nicotine exposure activates xenobiotic enzymes, which further stimulates feeding (23). Thus, perhaps unexpectedly, increased metabolic detoxification may promote rather than preclude toxin accumulation via increased feeding.

Intriguingly, while nicotine consumption increased A4 fly survival against parasitism, A4 flies under all but the nicotine-fed, unparasitized condition had lower viability than A3 flies (Fig. 2A). Thus, in a hypothetical population made only of A3 and A4 flies and exposed to *L. heterotoma* and nicotine, natural selection may be unlikely to favor A4 individuals. In this scenario, the evolutionary outcome would depend partly on whether antagonistic pleiotropy exists among loci determining metabolic resistance and viability. One general viability QTL has been identified in DSPR strains, but this QTL does not contain detoxification genes. Moreover, A4 and A3 flies seem to share the same allele at this QTL (15). Furthermore, while A4 survival was generally lower than A3, A3 (and not A4) female body mass was reduced by nicotine consumption. Body mass is correlated with fecundity in *D. melanogaster*, and thus nicotine-fed A4 flies may have greater reproductive success than A3 (24), which would potentially offset the cost of lower survival.

To our knowledge, *D. melanogaster* does not possess active nicotine sequestration mechanisms. Some drosophilids, such as *D. sechellia*, are known to acquire chemical defenses from toxic food sources (25), and *D. melanogaster* self-medicates against parasitoids using ethanol (26). However, other drosophilids that consume toxins have not been evaluated for chemical defenses (15, 16). Our finding that flies can utilize nicotine for defense without active sequestration mechanisms suggests that other organisms that tolerate toxin consumption could receive a transient defensive advantage, too. The biochemical properties and metabolic context of each toxin should affect their propensity to bioaccumulate. For example, non-toxic glucosinolates (GLS) rapidly breakdown into toxic mustard oils; thus, GLS-sequestration requires adaptations that interrupt this process (16). Many organisms sequester toxic steroids or alkaloids (4, 27, 28), perhaps because these more readily diffuse or are transported across tissues. Here we find that in addition to having increased nicotine metabolism, A4 *D. melanogaster* flies also likely consume much higher quantities of nicotine than A3 flies (Fig. 2C). We hypothesize that higher intake may allow relatively more nicotine to escape metabolism and permeate into the hemolymph of A4 flies, affecting *L. heterotoma* development to a greater degree than in A3 flies. This pattern could be verified with future studies that compare nicotine abundance in different tissues of A4 and A3 flies.

In conclusion, we find that elevated resistance increases passive toxin accumulation. Further, this accumulation produces a toxin-mediated fitness advantage against natural enemies, in animals without identified sequestration mechanisms. Reliance on metabolic detoxification is likely the ancestral character state for organisms with acquired chemical defenses, and variation in toxin metabolism is common (29). We therefore propose that one of the first steps in the evolution of chemical defense may paradoxically be natural selection for increased toxin metabolism.

## Methods

### Fly and wasp stocks

Flies were maintained at room temperature on molasses media from the Fly Food Facility at UCB; survival and parasitism experiments used Ward’s Instant Drosophila media to facilitate toxin dosing.

Wasps were maintained at room temperature on W118 *D. melanogaster* and 70%-honey water. Experiments used wasps within two weeks of eclosion.

### Generation of fly larvae

Approximately one-thousand flies were allowed to lay eggs for three days in three replicate resealable plastic containers with a layer of molasses-agar smeared with yeast paste. Larvae were hen pooled from each container, and second-instar larvae (L2) were selected based on morphology under a dissection microscope. Flies were not sorted by sex.

### Nicotine-resistance experiment

Twenty A4 and A3 L2 larvae were transferred one-by-one from egg-laying chambers into 5 replicate vials containing the following nicotine concentrations: 0 mM, 0.5 mM, 1.25 mM, 1.75 mM, 2.25 mM, 2.50 mM, 3.00 mM, 4.00 mM, 5.00 mM nicotine-treated media. Vials were checked daily for new pupae and eclosed flies, and daily counts were used to calculate developmental rate across nicotine doses (Fig. S1).

### Parasitism experiment

For each fly strain, 400 L2 were transferred into six replicate plastic containers containing molasses agar. Forty female and twenty male wasps were added to three containers (“wasp” treatment) while the other three were left unmanipulated (“no-wasp” treatment); all containers were left for 24hr. One “no-wasp” container contained only 80 L2s. The L2s were then counted individually (to avoid batch bias) into forty vials containing either control or 1.25-mM nicotine media. We pooled data on A4 flies from two separate runs of this experiment (average survival was not significantly different between runs). In run 1 (A4 only), we added 20 larvae to each vial. In run 2 (A4 and A3), we add 16 larvae to each vial. Vials were checked every 1-2 days for pupation and emergence. Parasitism was performed prior to nicotine treatment to avoid exposing *L. heterotoma* adults to nicotine. Therefore, changes in fly and wasp survival reflect the effects of nicotine consumption by *D. melanogaster* larvae and not any behavioral change by *L. heterotoma*.

### Nicotine accumulation experiment

One-thousand A4/A3 L2 were distributed one-by-one from egg-laying chambers into five 1.25-mM nicotine-treated vials. At five developmental stages (3^rd^-instar larvae, day-1 pupa, day-3 pupae [A4 only], day-1 adult, day-3 adult), we collected five individuals and washed them individually in glass dissection wells with DI H_2_O. Pupae were removed from vials prior to eclosion to avoid contamination of the adult exoskeleton with nicotine. Individuals from each stage for each vial were pooled and frozen at -20°C.

Frozen flies were thawed and soaked with methanol (50 μL) at room temperature for 2-3 days to reach equilibrium. Crude methanolic extracts were transferred to limited volume autosampler vials and injected directly. Gas chromatographic-mass spectrometric conditions were as previously described (30); full details are given in the Supplementary Material.

### Body Mass Measurement

300 A3/A4 L2 were placed one-by-one from egg-laying chambers into twenty vials containing either control or 1.25-mM nicotine media. Upon pupation, individuals were removed and placed onto food-free vials. Adults were starved for 48 hours and then weighed.

### Statistical Analysis

Statistical analyses were conducted using Rv3.6.1 (31). LC50s were calculated using adapted version of the ‘dose.p’ function from the ‘MASS’ package (32) to a binomial regression model of normalized percent survival versus nicotine dose generated by the ‘glmer’ function from lme4. Fly survival and wasp success were assessed by applying a least-squared-means test to a binomial regression model of survival as a function of nicotine and (for flies) parasite treatments using the ‘glm’ function from lme4 (33). Adult fly mass was compared by applying the least-squared-means method described above to a model of average mass per vial as a function of nicotine and sex. Developmental rate and mean nicotine content of flies was compared across strains using Wilcoxon signed-rank tests in base R.

### Data accessibility

Raw data files, R script, and detailed metadata are available for download from the Dryad Digital Repository: https://doi.org/10.5061/dryad.w3r2280sc (34).

## Supporting information

Metadata for Dryad files

## Acknowledgements

We thank Kirsten Verster, Jessica Aguilar, Mariana Karageorgi, Luis Jazo, Max Lambert for input on the experimental design, Noah Whiteman for helpful discussions, Stuart Macdonald for DSPR lines, Todd Schlenke for wasps. RDT was supported by the Miller Institute for Basic Research in Science, UC Berkeley start-up funds, and the Hellman Fellows Program; RWF was supported by NSF CCLI-DUE-0942345 and DEB-1556982.

---

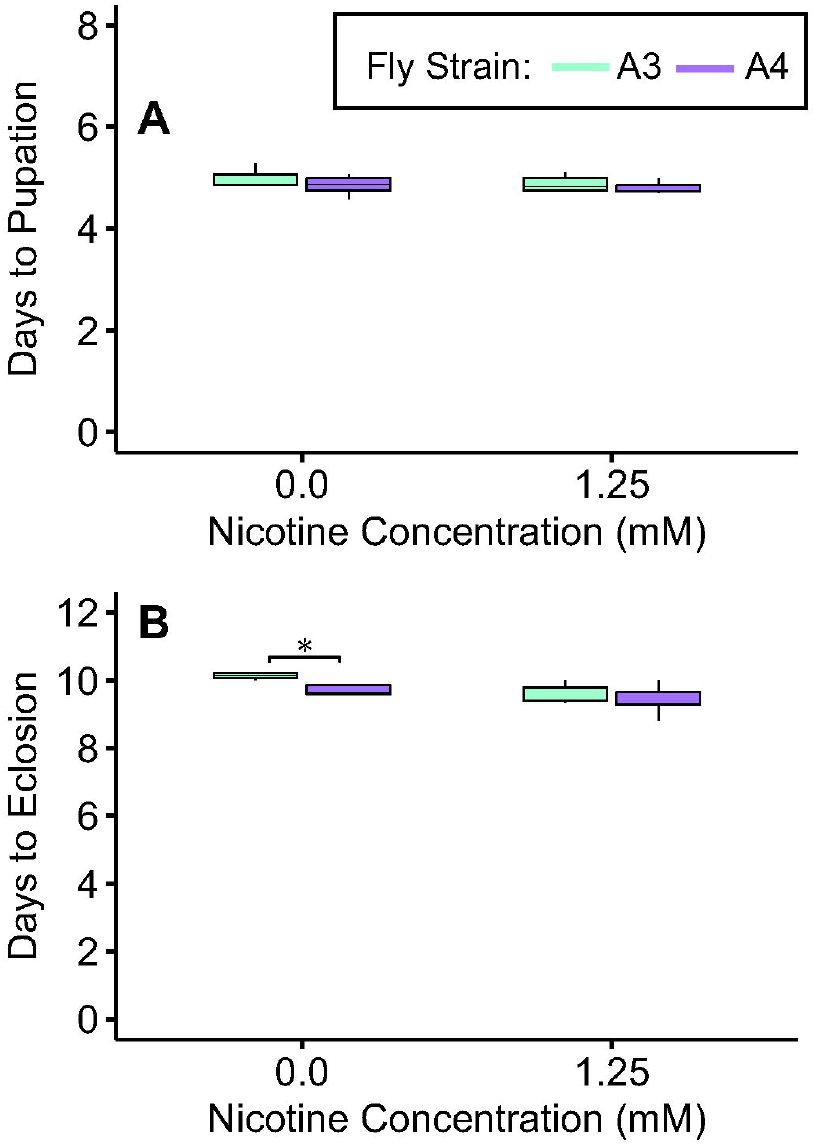

## Notes

https://doi.org/10.5061/dryad.w3r2280sc

### Competing Interest Statement

The authors have declared no competing interest.

### Summary of Updates

We updated the calculation of cotinine accumulated by each fly using a new method that allowed us to quantify the amounts in nanograms. Figure 2 has been updated accordingly. In addition, we revised the manuscript following requests from three anonymous reviewers. The most notable change is the addition of a discussion of whether A4 flies would be favored by selection given their overall low survival compared to A3 flies (see paragraph starting on line 147).

