## Supplementary Online Material for "Trade-offs between cost of ingestion and rate of intake drive defensive toxin use"

^+^ Authors listed in alphabetical order

*Corresponding authors:

Tyler Douglas

Rebecca Tarvin

3101 Valley Life Sciences Building

Berkeley, CA, 94720

**Supplementary Information:**

**Figure S1. A)** A3 and A4 time to pupation is similar on control and 1.25-mM nicotine food. **B)** A3 adults eclose significantly slower than A4 on control food, while days to eclosion is similar on 1.25-mM nicotine.

**GC-MS Analysis of samples.**

Frozen flies were thawed and soaked with methanol (50 μL) at room temperature for 48 hours. Crude methanolic extracts were transferred to limited volume autosampler vials and injected into a Thermo Trace GC, equipped with a Restek RTX-5MS 30 m x 0.25 mm capillary column 0.5 μL with He carrier at 40 cm/s, temperature programmed at 100°C held 1 min, then a 10°C/min ramp to 280°C and held 10 min. Samples (1 μL) were injected splitless at 250°C with a closed time of 1 min and surge pressure of 200 kPa. The GC was interfaced to a Thermo iTQ ion-trap mass spectrometer with a 250°C EI source, 70 eV ionization and autotuned with perfluorotributylamine, using automatic gain control (max ion time 25 ms). A solvent delay of 3 minutes was used. Nicotine di-dartrate (Sigma) standards were run before, midway through and at the end of test samples at log serial dilutions from 1.00 nM to 100 μM, eluting at 8.8 minutes and followed with blank methanol injections to avoid carryover. Integration analysis was done using extracted ion chromatograms at *m/z* 84 for selectivity (detection limit estimated at 0.2 ng (3 σ). The nicotine metabolite cotinine (eluting at 13.3 minutes) was identified by searching its mass spectrum. Later injection of an authentic nicotine and cotinine sample along with a representative fly extract confirmed the retention time and provided positive ID for cotinine. In the absence of quantitative standards for cotinine at the time of first analysis, we estimated relative rates of nicotine breakdown by ratioing the integrated area of cotinine at *m/z* 98 vs nicotine at *m/z* 84. Follow up analysis of nicotine and cotinine calibration curves showed a consistent response ratio of 2.97 for nicotine relative to cotinine. This allowed us to retroactively calculate the cotinine amounts in the samples.

Data were reviewed for but did not indicate presence of nicotine metabolites nornicotine and myosmine. Two other possible metabolites were found but remain unidentified. One nicotine-fed A3 pupal sample contained an order of magnitude more nicotine than all other measurements from both fly strains. Contamination by nicotine-treated media was likely the source of this inflated measurement, as the sample contained an extreme surplus of nicotine only and not its corresponding metabolic by-product, cotinine. We therefore excluded this outlier sample from our analysis. Due to concern over potentially low survival of A3 individuals on nicotine-treated media, we sacrificed individuals from only one pupal stage (day-1 pupae) to ensure that a sufficient number of survivors remained for sampling during adult stages. By contrast, we sampled two pupal stages from nicotine-resistant A4 flies.

Nicotine reference chromatogram at 10uM (10 pmol injected)

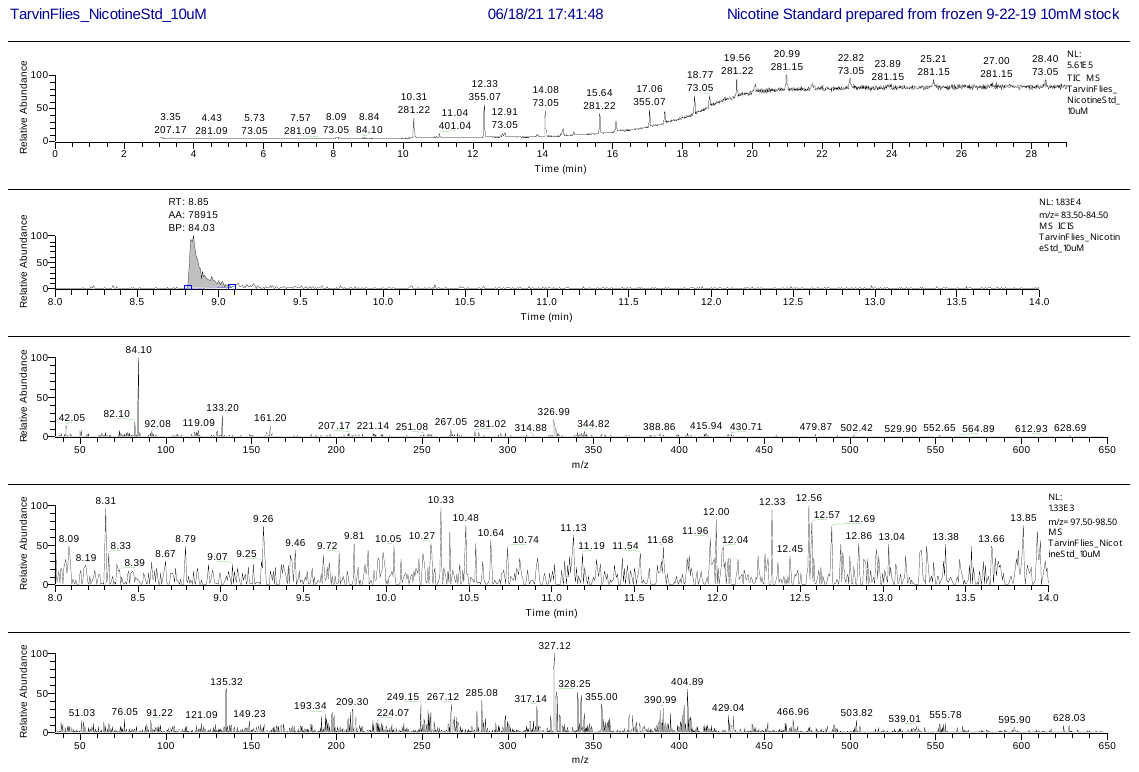

**Nicotine (EIC, *m/z* 84)**

**Total Ion Chromatogram**

Crude fly extract, A3 pupae (5 specimens in 50 uL methanol, 1 uL crude extract injected)

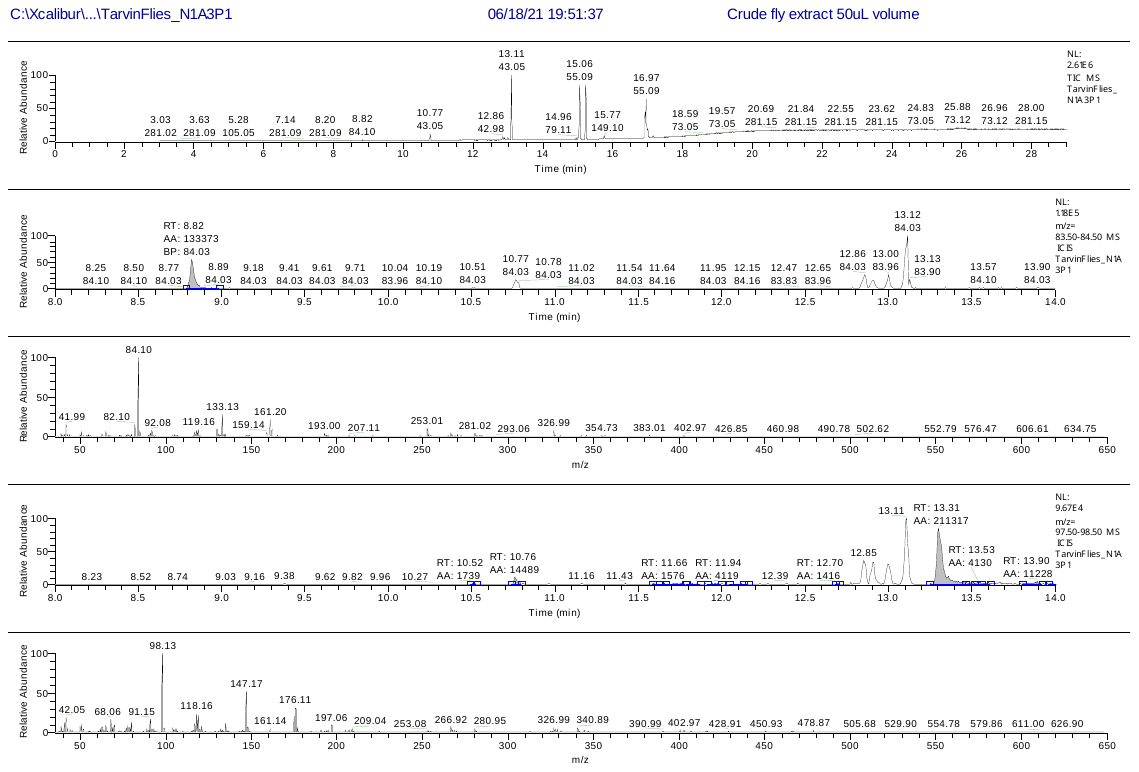

**Total Ion Chromatogram**

**Cotinine (EIC, *m/z* 98)**

**Nicotine (EIC, *m/z* 84)**

Calculation of relative response factors for cotinine to nicotine. Concentrations used reflect areas found in fly samples.

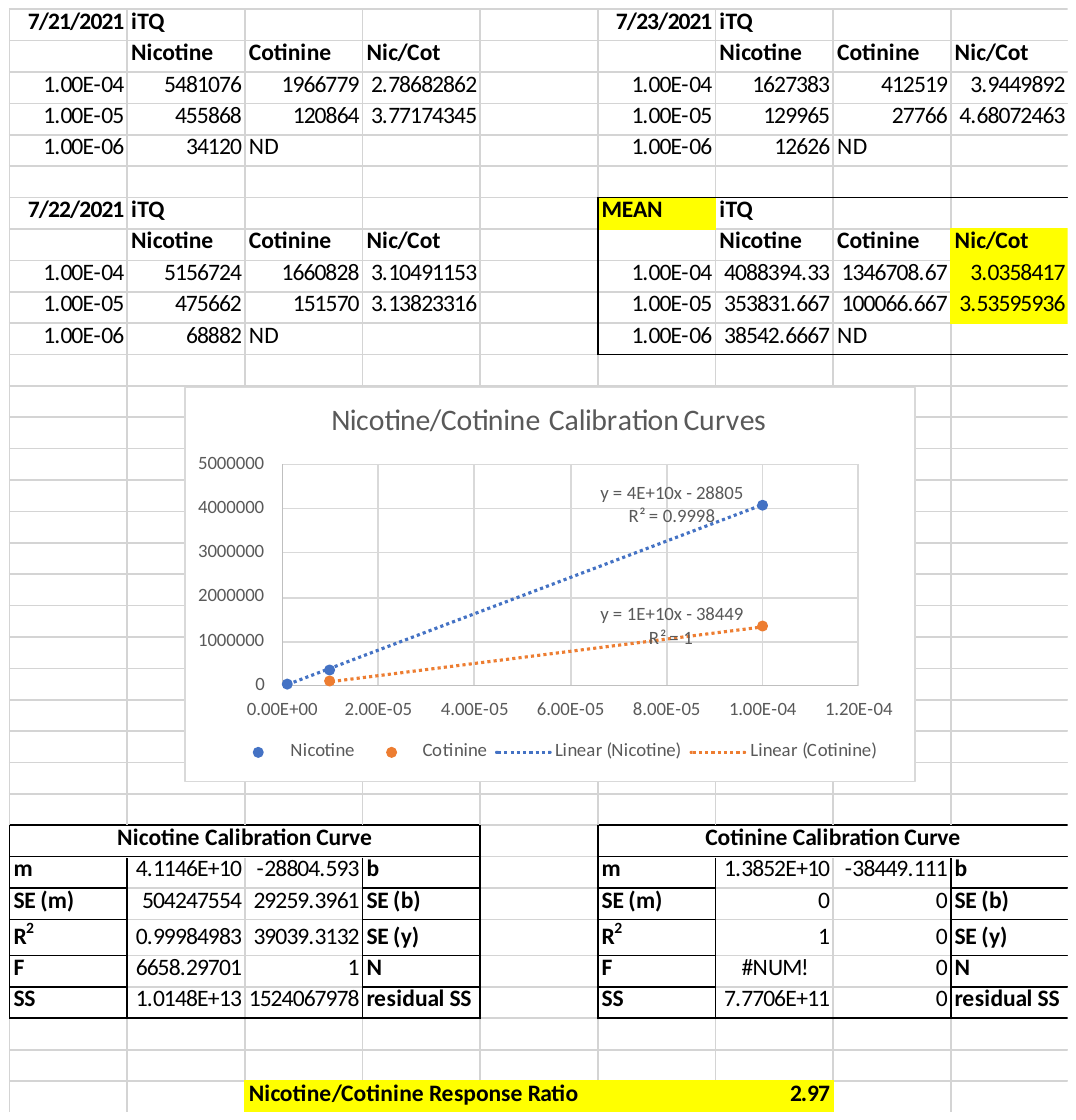
