## Supplementary material for "Trade-offs between cost of ingestion and rate of intake drive defensive toxin use": Metadata for Dryad files

^+^ Authors listed in alphabetical order

*Corresponding authors:

Tyler Douglas

Rebecca Tarvin

3101 Valley Life Sciences Building

Berkeley, CA, 94720

**Overview:**

All data were collected directly by contributing authors as described in the main text. All statistical analyses were completed using R version 4.1.1 and described in the main text. All statistical analyses and codes for figures are contained within a single annotated R script titled “Douglas_etal2021_data_analysis.R”. Raw data is stored in four csv files generated in Microsoft Excel (file details and column headers described below). We provide our data under the Creative Commons Attribution Non-commercial ShareAlike license (https://creativecommons.org/licenses/by-nc-sa/2.0/).

**Data accessibility:**

Raw data files and R script are available on Dryad at https://datadryad.org/stash/share/LW3IOIf2XbHqBEybyy-7RbcylCOuqzWg8QucvxUZWms

**Nicotine resistance data (Figure 1):**

Raw data of A4 and A3 nicotine resistance is stored in the file titled “nicotine_media_fly_survival.csv”. Each row of the datafile represents survival data from a single vial of *Drosophila* melanogaster. The columns titled strain, replicate, and dose indicate each vial's fly strain, nicotine concentration (in mM), and replicate number. The alive column indicates the number of flies that survived nicotine exposure out of 20 initial individuals, and the dead column indicates how many died. The percent column is percent survival (alive/20 *100).

**Parasitism survival data (Figure 2AB):**

Raw data of A4 and A3 parasitism survival is stored in the file titled “parasitism_survival.csv”. Each row represents survival data from a single vial. Strain, dose, and vial columns indicate fly strain, nicotine concentration (in mM), and replicate number as in “nicotine_media_fly_survival.csv”. A4 survival data were pooled between two independent runs of this experiment, and the exp_run column indicates whether a given vial comes from run 1 or 2. The alive and dead columns indicate fly survival/mortality out of 16 (in run 1) or 20 (in run 2) initial larvae, and raw_survival indicates raw survival as a percentage (alive/initial larvae*100). The norm_survival column is raw survival data divided by the highest individual survival value from each fly strain in the control, non-parasitized condition. The columns wasp_alive, wasp_dead, wasp_percent, and wasp_norm columns contain data parallel to the four fly columns described above but pertain to wasp success/failure.

**Body mass data (Figure 2C):**

Adult fly body mass data is stored in the file titled “adult_body_mass.csv”. Each row represents a single fly. The vial, strain, and treatment columns respectively indicate replicate, fly strain, and nicotine treatment (control = 0, nicotine = 1.25 mM). The weight_mg column contains individual fly mass.

**GCMS data (Figure 2D):**

GCMS data is stored in the file titled “GC-MS_Nicotine_Cotinine_flies.csv”. Each row represents a combined measurement of five flies obtained from the same vial of fly media, which were pooled together into a single GCMS sample. The label column indicates the sample name (i.e., replicate). The strain, stage, and treatment columns indicate fly strain, developmental stage, and nicotine media concentration (control = 0, nicotine = 1.25 mM), respectively. The columns “Nic_Area_perfly” and “Cot_Area_perfly” contain nicotine and cotinine GCMS integrated areas divided by 5 to represent the average amount per fly. The “Conc (M)” column contains the nicotine molarity of each GCMS sample. The “nicotine_ng/fly” and “cotinine_ng/fly” columns indicates the nanograms of nicotine and cotinine contained in each fly calculated based on molarity and molar mass. The “cot_nic_ng_ratio” column contains the cotinine area divided by the nicotine area of each fly.

**Adult developmental rate (Figure S1B):**

Adult developmental rate data is stored in the file titled “adult_developmental_rate.csv”. Each row represents a count of the number of adults eclosed in a given vial on a given day. The vial column contains information on fly strain, replicate, and nicotine treatment. The day column indicates the number of days since the start of the experiment. The adults column indicates the number of adults that eclosed in a vial on a particular day.

**Time to pupation (Figure S1A):**

Data on the time to pupation is stored in the file titled “pupal_developmental_rate.csv”. The data is formatted in the same manner as “adult_developmental_rate.csv” as specified above.

**References:**

1. R Core Team. 2021. R: A language and environment for statistical computing. R Foundation for Statistical Computing, Vienna, Austria
